# Naught all zeros in sequence count data are the same

**DOI:** 10.1101/477794

**Authors:** Justin D. Silverman, Kimberly Roche, Sayan Mukherjee, Lawrence A. David

## Abstract

Genomic studies feature multivariate count data from high-throughput DNA sequencing experiments, which often contain many zero values. These zeros can cause artifacts for statistical analyses and multiple modeling approaches have been developed in response. Here, we apply common zero-handling models to gene-expression and microbiome datasets and show models disagree on average by 46% in terms of identifying the most differentially expressed sequences. Next, to rationally examine how different zero handling models behave, we developed a conceptual framework outlining four types of processes that may give rise to zero values in sequence count data. Last, we performed simulations to test how zero handling models behave in the presence of these different zero generating processes. Our simulations showed that simple count models are sufficient across multiple processes, even when the true underlying process is unknown. On the other hand, a common zero handling technique known as “zero-inflation” was only suitable under a zero generating process associated with an unlikely set of biological and experimental conditions. In concert, our work here suggests several specific guidelines for developing and choosing state-of-the-art models for analyzing sparse sequence count data.

## 1 Introduction

Many high-throughput DNA sequencing assays exhibit high sparsity, which often exceed 70% in microbiome, bulk-, and single-cell RNA-seq experiments [1, 2, 3, 4]. Such sparsity can be problematic for modeling [5, 3, 6, 7], as common numerical operations like logarithms or division are undefined when applied to zero. Empirical benchmarks also suggest that the frequency of zeroes in datasets can affect false discovery rates in analyses like differential gene expression [8, 9].

Multiple approaches have been proposed for tackling the modeling problems posed by zero values in sequence count data. A common approach for addressing numerical challenges associated with taking the logarithm or dividing by zero is to add a small positive value, or pseudo-count, to the entire dataset prior to analysis [10, 5]. A more sophisticated approach is to model all counts (including zero values) as arising due to random counting involving the Poisson, negative binomial, or multinomial distributions [11, 12, 13, 14, 15]. Often these methods perform inference on the statistical properties of the entire datasets rather than a single observed zero count. Still more complicated models permit greater flexibility in the modeling of zero values by layering secondary random processes on top of random count processes. Examples of such models include zero-inflated negative binomial models [16] and Poisson zero-inflated log-normal models [17].

Although an abundance of methods have been proposed for handling zeros, it remains unclear when certain approaches are to be preferred over others. Empirical benchmarks comparing sequence analysis software packages [8, 9] do not isolate the effects of zero handling relative to other modeling decisions such as how to filter samples or normalize read depths, which in turn precludes offering specific guidance as to which approaches to modeling zero values are more or less appropriate. A conceptual debate has also emerged around the appropriateness of zero-inflation, with some arguing it is unnecessary [18, 19, 17], while others have suggested it can account for the large number of zero values in single-cell RNA-seq data [20, 21, 22, 23, 24, 25, 16, 26, 27, 28, 4], bulk RNA-seq data [29, 30, 31, 32], and microbiome sequencing data [33, 34, 35, 36, 37, 38, 39, 40, 41, 42]. This debate reflects controversy as to what kinds of processes give rise to zero values in sequence count data [18, 19, 17, 20].

In this Perspective, we re-analyze published datasets to show that alternative methods of zero-handling can lead to different inference outcomes. To understand the origins of these differences, we introduce a categorization scheme for zero generating processes (ZGPs) in sequence count data. While the precise ZGPs that contributed to a given dataset are typically unknown, we can use simulation to examine if there exist zero-handling models that perform well across a range of different ZGPs. Overall, our analyses reveal minimal conceptual and analytical support for the use of zero-inflated models to handle zeros in sequencing datasets. Our results suggest that simpler models avoiding zero-inflation are preferable for most tasks.

## 2 Real Data Examples

To investigate whether different methods of modeling zero values can affect the outcomes of real-world analyses, we reanalyzed six previously published datasets using models that differed only in their handling of zero values. We chose datasets that spanned a range of sequencing tasks: single cell RNA-seq [43, 44], bulk RNA-seq [45, 46], and 16S rRNA microbiota surveys [47, 48]. We then chose two different statistical models that differed only in their modeling of zero values. One model was based on a negative binomial distribution. Letting *y*_*ij*_ represent the observed counts for sequence *i* in sample *j*, a negative binomial model assumes that *y*_*ij*_ reflects the abundance of sequence *i* in sample *j* with added sampling noise described by a negative binomial distribution. Such negative binomial models are used in many popular software tools such as edgeR [49], DESeq2 [11]. The second model was similar to the first, but also assumed a process known as zero-inflation was taking place. In contrast to the first model, a *zero-inflated* negative binomial model additionally assumes that there exists a probability *π*_*j*_ that *y*_*ij*_ = 0 regardless of the abundance of sequence *i* in sample *j*. Zero inflation has become a popular method of augmenting the negative binomial to model higher levels of zero values in sequence count data [29, 20, 38, 16, 28]. To implement our two models, we used the ZINB-WaVE modeling framework [16], which allowed us to create identical negative binomial models that varied only according to the presence of zero inflation. Notably, our implementation relied on the default settings of ZINB-WaVE, which further assumes that the probability *π*_*j*_ can vary depending on the condition a sample belongs to (e.g. treatment or control). Such condition-specific zero-inflation is commonly used in a number of popular sofware packages [33, 20, 25, 16, 27]. We refer to these two models respectively as the Zero-Inflated Negative Binomial (ZINB) and Negative Binomial (NB) models (see Section 6 for more details).

To interpret the results of these two models we quantified the discrepancy between the top-K most differentially expressed sequences according to each model (Figure S1). Discrepancy was calculated as (*K* − *m*)/*K* where *m* is the number of the top-K sequences in common between the two models. We found that the ZINB and NB models disagreed on average by 46.3% (range: 20.0%-90.0%) among the top-50 most differentially expressed sequences. Even among the top-5 most differentially expressed sequences (a subset whose size and priority makes them likely for potentially costly experimental follow-up), disagreement averaged 50.0% and reached 100.0% for one dataset (Figure S1).

We found that the largest discrepancies between the ZINB and NB models occurred on sequences that were observed with a high number of counts in one condition, while also being observed with low or zero counts in the other condition (Figure S2). These presence-absence-like cases would seem like examples of where sequence abundance varies according to condition, and indeed, the NB model infers these sequences are differentially expressed (Figure 1). By contrast, we observed that the ZINB model does not always infer that sequences exhibiting presence-absence-like patterns in one condition were differentially expressed. The ZINB model instead inferred that these sequences were actually expressed at equal abundance; but, one condition exhibited higher rates of zero-inflation than the other condition (a phenomenon we term differential zero-inflation; Figure 1). Indeed, we found that there was a strong correlation between the difference in inferred differential expression according to the ZINB and NB models and the degree to which the ZINB model inferred that a sequence is differentially zero-inflated (Spearman rho > 0.35 and p-value ≈ 0 for all 6 datasets; Figure 1). Still, discrepancies between how the NB and ZINB models handled sequences with presence-absence like abundance patterns were not solely due to condition-specific zero-inflation, as we could recapitulate similar levels of discrepancy (average of 32.7%; range: 3.0%-48.0%; Figures S3-S5) even when condition-specific zero-inflation was disabled (see Section 6 and Supplemental Materials for complete discussion and methods). Ultimately, while the true processes underlying sparsity in a genomic dataset are often unknown, we found it striking that sequences with high counts in one condition, while also being observed with low or zero counts in the other condition, were often inferred by the ZINB model to not be differentially expressed. This suggest that zero-inflated models lead to higher false-negative rates than identical non-zero-inflated models.

**Figure 1:**
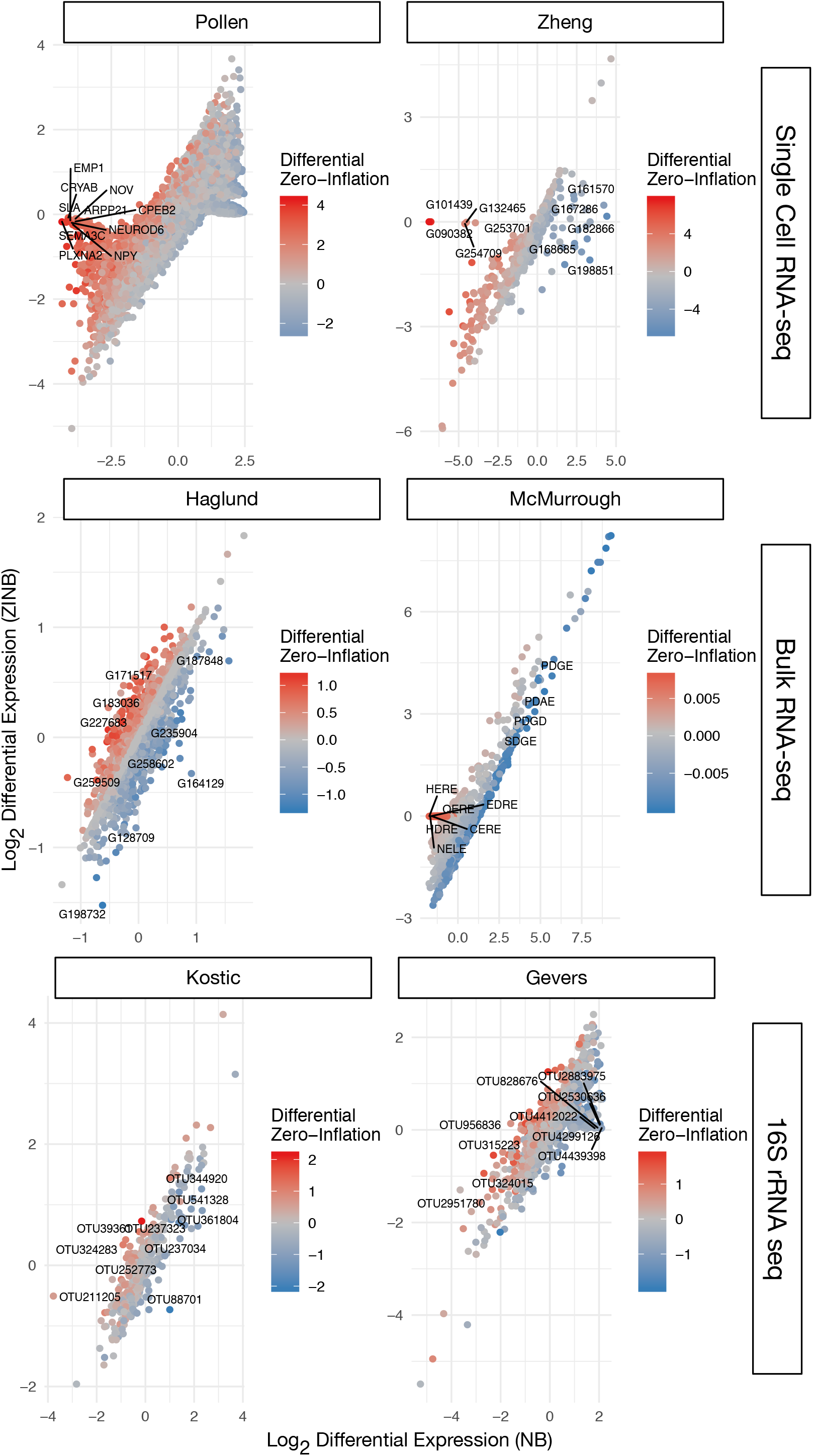
Differential expression (DE) estimates from a negative binomial (NB) and zero-inflated negative binomial (ZINB) model can differ substantially. Log base 2 differential expression for the ZINB and NB models are shown after each was applied to single cell RNA-seq, bulk RNA-seq, and 16S rRNA microbiota data. Dots represent different sequences, and each is colored according to the degree of differential zero-inflation as estimated by the ZINB model. For each dataset, the 10 sequences that have the largest discrepancy between inferred DE are labeled and their distribution in each condition is given in Figure S2.

## 3 Zero Generating Processes (ZGPs)

To provide a conceptual framework for analyzing how different zero handling models behave, we developed a scheme for categorizing different zero generating processes (ZGPs). Our scheme partitions ZGPs into three major classes (Figure 2):

**Figure 2:**
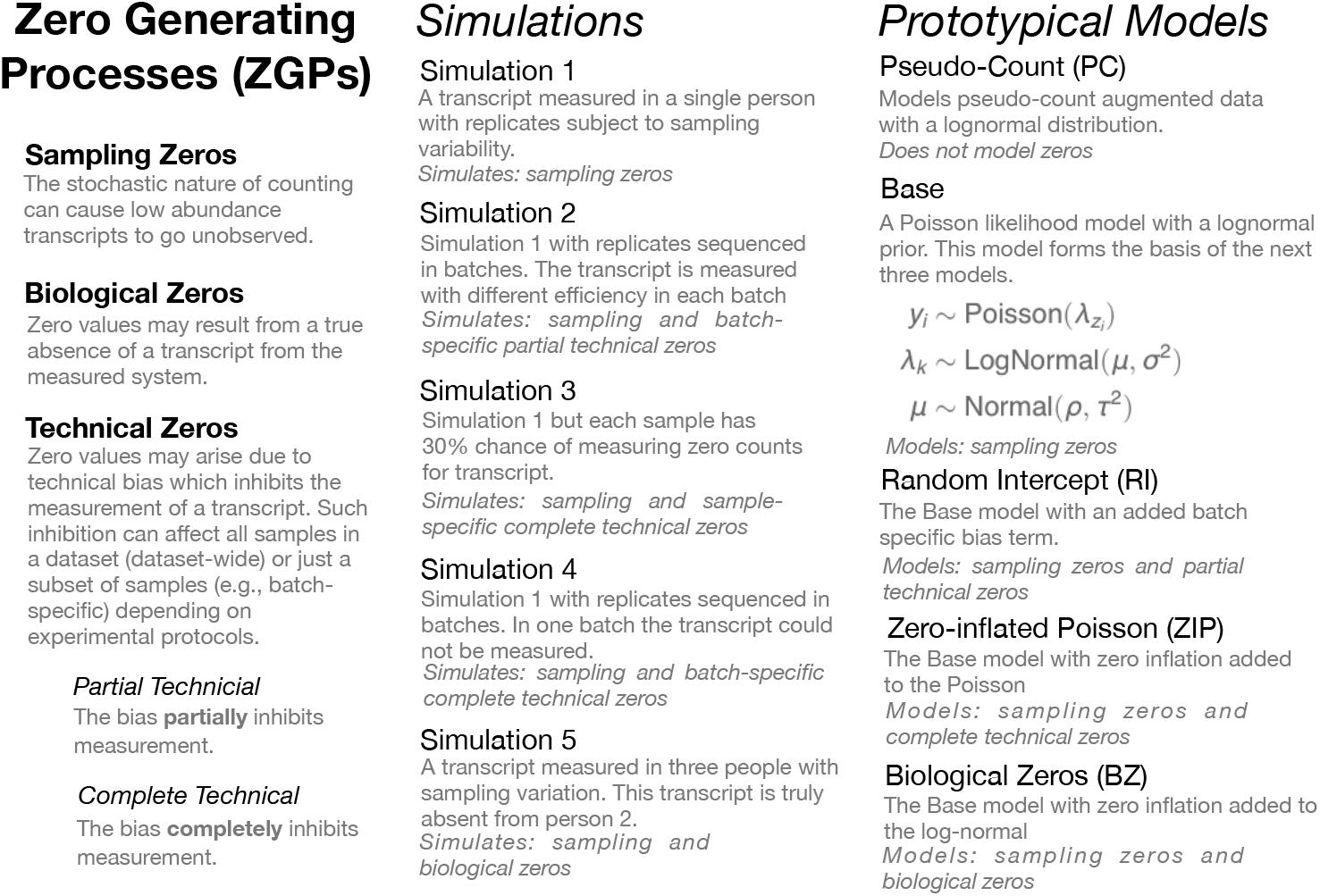
An overview of the zero generating processes (ZGPs), simulations, and models presented in this work. Model notation is as follows: *y*_*i*_ represents the number of counts observed for a given sequence in sample *i*, *z*_*i*_ represents the person from which sample *i* originates, *x*_*i*_ represents the batch number of sample *i*, 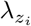 represents the abundance of the sequence in person *z*_*i*_. *σ*^2^, *ρ*, and *τ*^2^ are fixed hyper-parameters of the model.

### Sampling Zeros

Zeros may also arise due to limits in the total number of sequencing reads counted in a given sample [3]. Certain sequences, particularly ones at low abundance, may be present but not counted. In the limit where no reads were collected for a given sample, all zeros would be due to sampling effects; by contrast, if an infinite number of reads could be collected for a sample, sampling zeros would not be present.

### Biological Zeros

Perhaps the most intuitive reason for zeros in a dataset, biological zeros arise when a sequence is truly absent from a biological system.

### Technical Zeros

Preparing a sample for sequencing can introduce technical zeros into the data by ****partially**** or ****completely**** reducing the amount of countable sequences. These processes can lead to reductions in sequence abundance across all samples in a study or can act on a subset samples. For example, some genes are under-represented (partially reduced) in sequencing libraries due to the relative difficulty of amplifying GC-rich sequences [50, 51]. This bias could occur across all samples in a study; or, in a heterogeneous manner if different batches of samples were amplified using different primers or cycle number. Furthermore, if instead of a relative inability to amplify a sequence there was a *complete* inability, we would have a complete technical process. Zero-inflated models consider a specific case of complete technical zeros where this inability to measure a given sequence occurs randomly from sample-to-sample. Importantly, in a complete technical process, even abundant sequences may go unobserved.

There is ample experimental evidence that biological, sampling, and partial technical zeros occur in real data. Biological zeros are known to occur when studying gut microbiota across people [52] since unrelated individuals will harbor unique bacterial strains. Another example of biological zeros can be found in RNA expression analyses of gene knockout experiments, where gene deletion will eliminate certain transcripts from the expressed pool of genes prior to DNA sequencing [53]. Sampling zeros are known to occur when sequencing depth is limited and sequence diversity is high [2, 18]. For example, *in silico* studies have shown that decreasing the depth of sequencing studies can increase the numbers of observed zeros [36]. There also exist well-known examples of partial technical processes such as DNA extraction bias [54], batch effects [55, 56] or PCR bias [51, 57, 58, 59].

In contrast with the other three ZGPs, the rationale for modeling complete technical zeroes is more nuanced. Dataset-wide complete technical zeros certainly exist: certain prokaryotic taxa are known to not be detectable by common 16S rRNA primers, for example, and will therefore not appear in microbiota surveys [60, 61]. Of course, modeling the expression of dataset-wide technical zeros is typically neither considered a useful nor practical endeavor. Much more interest and effort though has been invested in considering cases of sample-specific complete technical zeros; that is, when with some probability *π*_*j*_, a sequence *j* may go completely unobserved in a given sample, regardless of its true abundance. Such sample-specific complete technical zeros have been considered in the analysis of gene-expression data, where zero-inflation has previously be used to model a variety of processes often termed “dropout” [20, 22, 23, 25, 27]. Dropout has been explained as stochastic forces involved with sampling of low-abundance sequences or due to the stochastic nature of gene expression at the single-cell level [20, 22]. Dropout has also been described as the result of failures in amplification during the reverse-transcription step in RNA-seq experiments [20, 22]. In the analysis of microbiome data, zero-inflation has been used to model differing presence/absence patterns between individuals [20, 22]. Yet, each of the aforementioned phenomena could be argued as arising from either sampling, biological, or partial technical ZGPs. The inability to detect low abundance RNA sequences could be argued as arising from a sampling process rather than a complete technical process. Stochastic gene expression at the single-cell level and differing presence/absence patterns between individuals fit the definition of biological zeros. Last, zeros associated with gene amplification inefficiency could either reflect a sampling process or a partial technical process related to competitive inhibition between sequences in the amplification reaction. Thus, models considering sample-specific complete technical processes may actually be attempting to capture other ZGPs.

## 4 Simulation Studies

As the ZGPs present in a given dataset are typically unknown, we sought to understand how different methods of handling zeros performed under model mis-specification. Exploring model behavior using empirical benchmarks that compare common software platforms (e.g. DESeq2, EdgeR, or ALDEx2) is not optimal because multiple modeling decisions independent of zero-handling like data filtering, inference method, and data normalization are incorporated into commonly used sequence analysis tools. To isolate the effects of different zero-handling methods on sequence analysis, we therefore created a simple inference model based on the Poisson distribution (Base model, Figure 2). We chose this distribution in place of the negative binomial (also called the Poisson-Gamma) distribution used in our earlier analyses because the Poisson yields simpler to understand, single-parameter inference, while still modeling random sampling.

We then layered four common zero-modeling approaches onto our Base model (Figure 2; full descriptions of each model are presented in *Methods*). We created a Zero-Inflated Poisson (ZIP) model by adding a zero-inflation component to the Poisson distribution in the Base model. To model zeros arising due to true biological absence, we created the Biological Zero model (BZ). This model includes a zero inflated component on the Log-Normal portion of the Base model and is similar to the DESCEND model of Wang et al. [17]. To model sampling and partial technical zeros, we created the Random Intercept (RI) model. The RI model differs from the Base model only by allowing external covariate information, *e.g.*, batch numbers, to decrease the amount of available sequences. Last, we designed a Pseudo-Count (PC) model that does not model any ZGPs, but rather avoids numerical issues with zeros by adding a fixed positive pseudo-count *κ* to each observed count value (i.e., each cell) in the data before analysis. The PC model removes the Poisson component from the Base model and instead uses 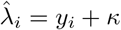 as the observation.

We next designed five different simulation experiments to test our hypothesis that some zero-handling methods would be more tolerant to model mis-specification than others. As sampling lies at the heart of sequence count data [62, 18], we included zeros from Poisson sampling in each simulation. In addition, to investigate the other ZGPs, the second through fifth simulations included biological, complete-, and partial-technical processes as well. Each model was applied when simulations could distinguish models. For example, in the absence of covariate information the RI and Base models are identical; therefore, the RI model was only applied to simulations where covariate information was available and could distinguish the RI and Base models (*See Supplementary Materials for a full discussion of this topic*).

We judged model performance on a given simulation based on the inferred probability that the sequence’s abundance was less than or equal to the true simulated value, *i.e.*, the cumulative distribution function of the posterior density evaluated at the true value. This statistic captures both the error in the model’s best guess (the mean) as well as its certainty about that guess (the spread of the posterior about the mean; Figure S6 provides a visual explanation of this metric). An optimal model would have a value of 0.5, whereas models that performed poorly would have values near 0 or 1 if they under- or over-estimated the true value with undue certainty.

The results of our simulation studies are shown in Figure 3 and S4. (A detailed description and explanation of the results is presented in *Supplementary Materials*.) Overall, we found the best-performing model to be the RI model. In particular, this model displayed three beneficial features. First, like the Base model, the RI model appropriately modeled sampling zeros without difficulty. Second, even though it did not model biological zeroes directly, the RI model approximated biological zeros as very low abundance; conveying the key information, that the absent sequence was not common. Third, by allowing covariate adjustment, the RI model effectively estimated sequence abundance even in the presence of partial technical zeros due to batch effects.

**Figure 3:**
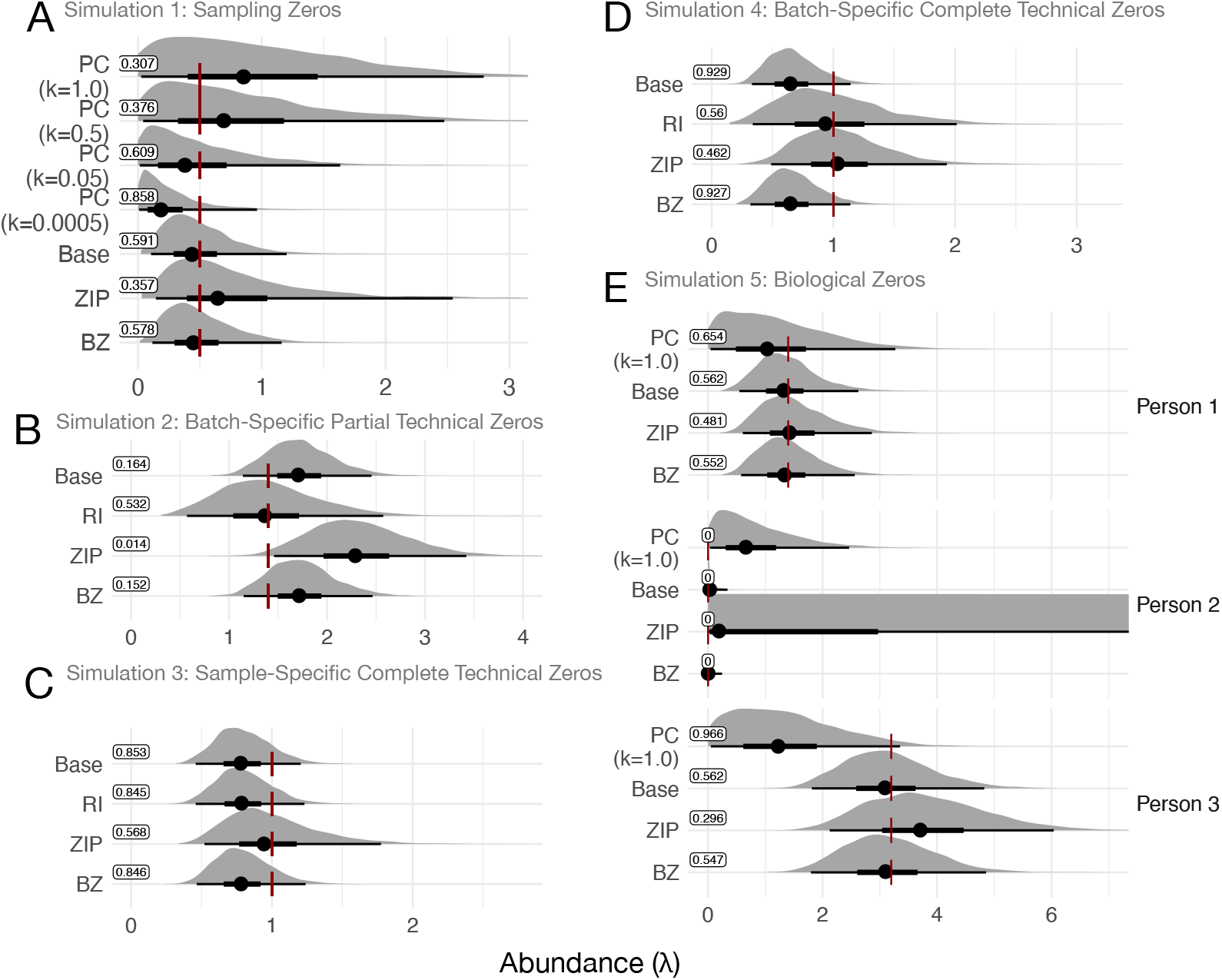
A summary of how different zero handling models behave on different simulations of zero generating processes. Shown are posterior distributions of sequence abundance (*λ*) from each model. The dark red vertical bar represents true value of *λ*. Posterior mean as well as the 66% and 95% credible intervals are shown in black. Boxed values represent the cumulative distribution function of the true value of *λ*, as described in the text and in Figure S6, the best performance possible is a value of 0.5. The statistic is by definition zero when *λ^true^* = 0 (*e.g.*, person 2 in panel E) in which case performance is assessed visually. (A) Simulation 1 (sampling zeros only), (B) Simulation 2 (batch-specific partial technical and sampling zeros), (C) Simulation 3 (sample-specific complete technical and sampling zeros) (D) Simulation 4 (batch-specific complete technical and sampling zeros), (E) Simulation 5 (biological and sampling zeros). In panel E, the abundance axis (*λ*) was cropped to enable all model results to be shown.

The next best performing models were the Base and BZ models. The Base model exhibited similar simulation results as the RI model, with the exception of its performance on technical zero simulations due to its inability to incorporate additional covariates like batch. The BZ exhibited a similar limitation on modeling external covariates and hence, batch effects. Still, the BZ model performed well on modeling of biological zeroes, which was expected because it is designed specifically for such a phenomenon.

More poorly performing models in our simulations were the PC and ZIP models. In addition to its observed lower accuracy, the PC model was also sensitive to chosen pseudo-count values (Figure 3A and E). The ZIP model overestimated sequence abundances (*λ*) in every simulations with the exception of the one explicitly simulating sample-specific complete technical zeroes (Simulation 3). Additionally, our simulations showed that the ZIP model had high posterior uncertainty: The ZIP model could not tell if zeros were due to sampling of a low abundance sequence, technical censoring of a low abundance sequence, or technical censoring of a high abundance sequence (Figure S8). Such uncertainty biased parameter estimates even when more than 1000 replicate samples were available to the model (Figure S9). That is, even when the ZIP model had access to many replicate samples, the model falsely inferred that zero-inflation was present even when it was not. Our simulation of biological zeros also showed that, when used to model condition-specific zero-inflation like in the ZINB model of Section 2, posterior estimates of *λ* under the ZIP model can be the highest in conditions when true sequence abundances are the lowest (Person 2; Figure 3E). More broadly, our simulations recapitulate our earlier findings involving published datasets and provide examples for how zero-inflated models can spuriously infer sequences to be present even when unobserved.

## 5 Discussion

Here we have demonstrated, using real-world datasets, that different methods for modeling zeros can lead to disagreement among almost half of sequences when carrying out differential abundance analysis. We also categorized zero generating processes (ZGPs) and summarized evidence in favor of each. Last, we used simulations to explore how different zero handling methods perform when different ZGPs are present. In concert, these analyses suggest caution when considering zero-inflated models for handling zero values. These models may increase false negative rates in differential expression analyses, capture ZGPs that may actually be described by simpler biological processes, and are sensitive to model mis-specification.

Our conceptual framework and simulations provide insight into the mechanisms underlying the outcomes of empirical benchmarking studies [8, 9, 63, 42, 64]. Thorsen et al. [8] report that the zero-inflated Gaussian model metagenomeSeq increasingly biases estimates of differential abundance as data sparsity increases. Similarly, Dal Molin et al. [64] observe that models using zero-inflation, such as SCDE [20], or Monocle [65], have a higher false-negative rate for differential expression than alternative non-zero-inflated models such as DESeq [11] or edgeR [49]. Our results suggest these errors arise when zero-inflated models are applied to datasets where sample-specific complete technical processes are insubstantial.

Beyond explaining empirical phenomena, our framework suggests three guidelines for modeling sequence count data. First, for designers of new sequence count models, biological zeros can be approximated as sampling zeros. This recommendation is logical as biological zeros can be considered sampling zeros from a sequence whose abundance is vanishingly small. Moreover, our results demonstrate that sampling models will correctly interpret zeros as evidence of low-abundance sequence. Treating biological zeros as evidence of very low abundance sequence should encourage simplicity in future models, and is also consistent with the design of several existing tools such as Deseq2 [11], Aldex2 [66], MALLARD [13], Stray [67], GPMicrobiome [12] and MIMIX [14].

Our second guideline is that zero inflated models should be avoided. Our re-analysis of datasets from single cell RNA-seq, bulk RNA-seq, and 16S rRNA microbiome sequencing experiments suggested that zero-inflated models can result in the spurious conclusion that sequences are not differentially expressed when clear presence-absence patterns exist between experimental groups. Additionally, our literature review and categorization of ZGPs revealed that common motivating phenomena for zero-inflated models such as dropout can actually be considered forms of other common ZGPs. Last, we found that when sample-specific complete technical processes are not present in data, zero-inflated models produce biased estimates. Overall, this guideline is aligned with recent research demonstrating that after controlling for biological zeros in droplet single-cell RNA-seq experiments, zero-inflation is not necessary to describe the zero patterns observed in sequencing data [18]. Rather, zeros in these experiments were nearly perfectly captured by negative binomial models lacking zero-inflation (*i.e.*, without the need to model complete technical zeros). Thus, when sequence count data have more zeros than can be adequately modeled by a Poisson distribution, our guideline suggests that these “excess-zeros” can be modeled using sampling, partial technical, and biological processes.

Our third guideline for modeling zero values is to employ simple count models such as the Poisson, negative binomial, or multinomial that can incorporate technical covariates (e.g., batch). We underscore that this guideline does not require knowing the true ZGPs present in a study. Moreover, we do not expect that models will be able to reliably make this distinction either. Rather, our simulations show that simple models capable of accounting for both sampling and partial technical zeros can produce accurate inferences under a range of different ZGPs. Fortunately, such models have also already been implemented for a range of applications including generalized linear regression [67, 14], non-linear regression [67, 68], clustering [69], time-series analysis [13, 12, 67], and classification [70, 71].

Our guidelines for zero handling will eventually need to be incorporated into broader pipelines for the analysis of sequence count data. Other outstanding challenges exist in this arena. Tasks like gene and sequence variant calling or data processing can be considered to be independent of zero handling as they are often done as an isolated step prior to data modeling. Recent advances in addresses these independent tasks [72, 73, 74] should therefore combine in a straightforward manner with our zero handling guidelines. On the other hand, some tasks, such as data normalization, in sequence count data analysis are likely to require solutions that interact with a given zero handling framework. Data normalization choices, for example may convert integer counts into continuous variables [62] and thereby remove key information needed to understand sampling zeros. Future zero handling studies may therefore be combined with empirical studies of full sequence data analysis pipelines to provide additional insight into how modeling choices ultimately affect sequence count data analysis.

## 6 Methods

### 6.1 Analysis of Previously Published Data

We analyzed six previously published datasets. Each dataset was pre-processed such that only sequences observed in at least 3 samples with at least 3 counts were retained. For each dataset, the ZINB and NB models were fit using default parameter values, an intercept, and a binary variable denoting which of two groups samples belonged to. The NB model was created from the ZINB model by additionally specifying the matrix parameter *O_pi*. Large negative values for this parameter reduce zero inflation. The ZINB model used the default value for *O_pi* to allow zero inflation; the NB model set *O_pi* to be a matrix populated with the number −10^6^ to ensure no zero inflation was used. The non-condition-specific ZINB model was created from the ZINB model by additionally specifying which_X_pi = 1 and which_V_pi = 1. Details regarding how each dataset can be obtained, the groups compared, and the resulting dataset sparsity level are given in Supplementary File 1.

Differential zero-inflation was reported directly by ZINB-WaVE. The ZINB-WaVE model infers a zero-inflated parameter *π*_*j*_ on the logit scale. For the condition-specific ZINB model, *π*_*j*_ is described by the linear model 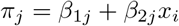 where *x*_*i*_ is condition of sample *x*_*i*_. Therefore, the paraeter *β*_2*j*_ can be interpreted as the degree to which the model differentially uses zero-inflation in one condition compared to another for sequence *j* – hence we termed this parameter differential zero-inflation.

### 6.2 Simulation Studies

First we introduce our notation.

*y*_*i*_ The number of counts of a specific sequence in the *i*^*th*^ sample
*z*_*i*_ The biological sample where sample *i* originates
*x*_*i*_ The batch in which sample *i* was processed

The following five models assume that each of the *K* biological specimens has a true parameter *λ*_*k*_ that represents the abundance of a single sequence *j*.

#### 6.2.1 Prototypical Models

For comparability, each of our five models are based on a hierarchical Poisson log-normal model. Letting *κ* denote a fixed non-zero value, we define the **pseudo-count (PC) model** as

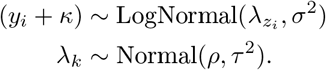

This model avoids numerical issues with taking the log of zero values by adding a pseudo-count to the data prior to analysis.

We defined the **base model** as

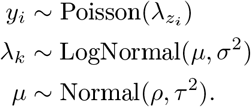

This model considers count variation and zeros due to sampling.

The **random intercept (RI) model** modifies the base model with a batch-specific multiplicative factor *η_x_i__*, which may alter the rate of Poisson sampling.

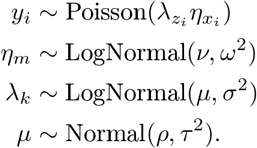

For identifiability, we assume that a single batch, labeled batch number 1, is an unbiased gold standard, so *η*_1_ = 1. If we have a single batch, this model is identical to the base model. The NB (negative binomial) model of Section 2 is similar to the RI model but uses the more flexible negative binomial distribution instead of the Poisson.

Like the ZINB model in Section 1, we created a **zero-inflated P oisson (ZIP) m odel** by adding a zero-inflated component to the Poisson part of the base model. The ZIP model is defined by

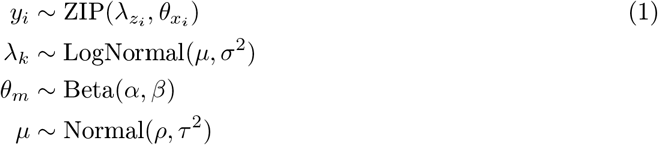

where 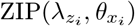 is shorthand for

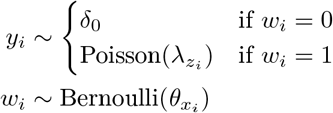

and where *δ*_0_ refers to the Dirac distribution centered at zero. This model assumes that all zeros arise due to a sampling process or a complete technical process.

In contrast to the ZIP model, the **Biological Zero (BZ) model** adds a zero-inflated component to the log-normal part of the base model. The BZ model is defined by

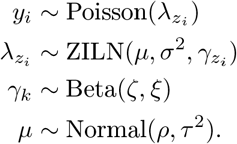

Here 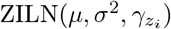 is short for:

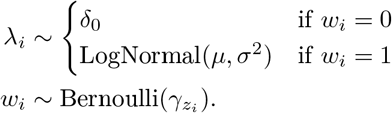

this model assumes that zeros arise from a sampling process or a biological process. In Section 4, the BZ model was modified from this form because of the difficulty of representing the latent Dirac distribution using the Hamiltonian Monte Carlo. Instead, the Dirac distribution in the BZ model was approximated with a truncated normal distribution with mean 0 and variance 0.0001.

#### 6.2.2 Simulations

Our series of simulation studies investigated the behavior of each model on each zero generating process. We present only univariate simulations. Hyper-parameter values were chosen to use in each of the five simulations. The hyper-parameters are: *σ*^2^ = 3, *ρ* = −1, *τ*^2^ = 5, *ν* = 0, *ω*^2^ = 2, *α* = .5, *β* = .5, *ζ* = 1, and *ξ* = 1. Simulations that had low likelihood under the simulating model were rerun. This procedure ensured that each simulated dataset contained enough information to recover the true parameter values. This was done for simulations in Figure 3 but not for simulations in Figure S9.

##### Simulation 1: Sampling Zeros

The first simulation consisted of 5 random draws from a Poisson distribution with a rate parameter *λ* of 0.5. This represents a single sequence within a single person, measured with 5 technical replicates all processed in the same batch. We applied the PC model with three different pseudo-counts: 1, .5, and .05.

##### Simulation 2: Sampling and Batch-Specific Partial Technical Zeros

The second simulation consisted of 15 replicates samples split into 3 batches with Poisson rate parameters 1.4, 0.6, and 3.2. This simulates polymerized chain reaction (PCR) efficiency varying by batch. As discussed above, batch 1 is derived from some gold standard measurement device with no bias.

##### Simulation 3: Sampling and Sample-Specific Complete Technical Zeros

The third simulation consisted of 15 replicate samples from a Poisson distribution with rate parameter *λ* of 1. This simulates a single sequence measured with technical replicates where each replicate has a 30% chance of catastrophic error, causing a complete inability to measure that sequence. This simulation was contrived to reflect the assumptions of the ZIP model.

##### Simulation 4: Sampling and Batch-Specific Complete Technical Zeros

This simulation uses sampling and complete technical zeros, and represents a single sequence measured in 15 replicate samples in 3 batches. However, because of a different reagent or missed experimental step, batch 2 complete lacked the sequence. We assume that no other bias is present in batches 1 or 3, which are represented as random draws from a Poisson distribution with rate parameter 1.

##### Simulation 5: Sampling and Biological Zeros

The fifth simulation consisted of 15 samples from three individuals with Poisson rate parameters 1.4, 0, and 3.2. This simulates the abundance of a single sequence measured in three individuals, of which two possess that sequence and one does not. To model biological zeros with zero inflation, we slightly modify Equation (1) in the ZIP model, replacing 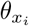 with 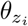. This change reflects a change of modeling zero-inflation by batch to modeling zero-inflation by individual. This corresponds to modeling condition-specific zero-inflation as in the ZINB-WaVE model.

#### 6.2.3 Posterior Inference

All 5 models were implemented in the Stan modeling language which uses Hamiltonian Monte Carlo (HMC) sampling [75]. Model inference was performed using 4 parallel chains, each with 1000 transitions for warmup and adaptation and 1000 iterations collected as posterior samples. Convergence of chains was determined by manual inspection of sampler trace plots and through inspection of the split 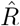 statistic.

### 6.3 Code availability

All code necessary to recreate the analysis and figures in this work is available at: https://github.com/jsilve24/zero_types_paper.

## Acknowledgements

We thank Rachel Silverman for her manuscript comments. JDS and LAD were supported in part by the Duke University Medical Scientist Training Program (GM007171), the Global Probiotics Council, a Searle Scholars Award, the Hartwell Foundation, an Alfred P. Sloan Research Fellowship, the Translational Research Institute through Cooperative Agreement NNX16AO69A, the Damon Runyon Cancer Research Foundation, the Hartwell Foundation, and NIH 1R01DK116187-01. SM and KR would like to acknowledge the support of grants NSF IIS-1546331, NSF DMS-1418261, NSF IIS-1320357, NSF DMS-1045153, and NSF DMS1613261.

## Author Contributions

JDS was responsible for conceptualization, methodology, data analysis, investigation, visualization, and writing. KR was responsible for writing. SM was responsible for supervision, writing, and funding acquisition. LAD was responsible for supervision, writing, and funding acquisition.

## Competing Interests

The authors declare that they have no competing financial, professional, or personal interests that might have influenced the performance or presentation of the work described in this manuscript.

## Supplement

### 1 Detailed Description of Simulation Results

Here we extend the discussion in the main text and give a detailed description of our simulation results. We want to intuitively explain why these results appear the way they do.

#### 1.1 Simulation 1: Highlighting Sampling Zeros

The first simulation consists of five random draws from a Poisson distribution with a rate parameter *λ* of 0.5. This simulation represents a single transcript within a single person measured with 5 technical replicates, all processed in the same batch. The small value of *λ* ensured that the data would contain sampling zeros with high probability. We applied the PC, Base, ZIP, and BZ models to this simulation. To demonstrate the impact of the choice of pseudo-count on the PC model, we applied the PC model with three different pseudo-counts: 1, .5, and .05. We summarize and provide an intuitive explanation of the results (shown in Figure 2A) below:

##### PC model

The PC model is sensitive to the choice of pseudo-count *κ*. Typical values for *κ* used in the analysis of sequence count data include .5, .65, and 1, as we cannot directly infer a generally optimal value from the observed data [1]. Here we found that *κ* = 0.05 provided a close correspondence between the posterior mean of *λ* and the true simulated value of *λ*.

##### Base model

The base model performs well, placing the posterior mean near the true simulated value of *λ*.

##### ZIP model

While the ZIP model is capable of modeling pure sampling zeros (*i.e.*, if *θ*_1_ = 0), this model substantial inflated *λ* compared to its true value. The ZIP cannot distinguish between zero values due to low abundance and low zero inflation (small *λ* and small *θ*) and zero values due to high abundance and high zero inflation (large *λ* and large *θ*). This interpretation is supported by a strong positive correlation in the posterior distribution of *λ* and *θ* shown in Figure S8. Figure S8 demonstrates that the regions of high posterior probability are spread out over a large range of possible *λ* and *θ* values. This uncertainty also appears in the long tails of the ZIP model’s posterior distribution for *λ*.

##### BZ model

The BZ model performs nearly identically to the base model. The presence of non-zero counts makes it extremely unlikely that the true value of *λ* is zero; if *λ* = 0 we would expect all counts to be zero. The BZ model estimates that the true value of *γ* must be near zero. If *γ* ≈ 0 then the BZ model reduces to the base model.

We repeated this analysis at a variety of sample size between 5 and 1280 with the same rate parameters as above. For each sample size we simulated 30 datasets. For each simulated dataset, we fit both the base and ZIP models. The distribution of the posterior means of each of these two models as a function of sample size is shown in Figure S9. With increased sample size, the inflation of *λ* decreases, but even with 1280 samples per dataset, the ZIP model continues to demonstrate inflation of mean estimate of *λ*. In contrast, with only 5-10 samples, the base Model estimates *λ* near its true value.^1^ Thus estimates from zero-inflated models can demonstrate bias even for extremely large sample sizes.

#### 1.2 Simulation 2: Highlighting Batch-Specifc Partial Technical Zeros

The second simulation consists of 15 replicates samples split evenly into 3 batches with Poisson rate parameters 1.4, 0.6, and 3.2. This simulation represents a situation where polymerized chain reaction (PCR) efficiency varies by batch. We consider batch 1 to be derived from some gold standard measurement device that has no bias. As the rate parameters for each batch are all small, this dataset contains a mix of sampling and partial technical zeros. We summarize and provide an intuitive explanation of the results (shown in Figure 2B):

##### Base model

The base model cannot incorporate batch information and therefore naively estimates that all 15 samples come from a distribution with a fixed rate parameter. The base model estimates the rate parameter as the mean of the rate parameters of the three batches. As this mean rate is higher than the batch 1 rate, the base model inflates its abundance estimate.

##### RI model

The RI model performs well in this simulation placing the posterior mean near the true value of *λ*.

##### ZIP model

The posterior mean of the ZIP model lies higher than that of the base or BZ models. This may seem surprising because the ZIP model can use batch information. This result can be understood in two parts. First, the ZIP cannot detect a shift in the overall Poisson rate parameter between batches; it can only detect differences in the rates of zeros between batches. This limitation causes the ZIP model to view the data and inflate estimates, like the base model does, based on the overall average rates between batches. Second, the zero inflation component of the ZIP model excludes some zero values from its estimates of *λ* and in doing so inflates the overall estimates for *λ*. Combining these two parts, the ZIP results can be seen as inflation as the base model does, plus more inflation due to its zero inflated component.

##### BZ model

Here the BZ model behaves identically to the base model. As in simulation 1, this occurs due to the presence of non-zero counts making it highly unlikely that *λ* = 0.

#### 1.3 Simulations 3 and 4: Highlighting Sample-Specific and Batch-Specific Complete Technical Zeros

The third simulation consists of 15 replicate samples from a Poisson distribution with rate parameter *λ* of 1. This simulation represents a hypothetical situation: a single transcript is measured with technical replicates; each replicate has a 30% chance of catastrophic error causing a complete inability to measure that transcript. As with prior simulations, the small rate parameter ensures that the data contains sampling zeros and complete technical zeros. We summarize and provide an intuitive explanation of the results (shown in Figure 2C):

##### Base model

The base model underestimates *λ*. The base model incorrectly assumes the complete technical zeros are really sampling zeros. The excess zeros thus deflate the base model’s estimates of *λ*.

##### RI model

Since all samples came from the same batch, there is no difference between the base and RI models. So the RI model also underestimates the true value of *λ*.

##### ZIP model

The ZIP model performs well, placing the posterior mean of *λ* near its true simulated value.

##### BZ model

As in simulations 1 and 2, the presence of non-zero counts makes it highly unlikely that the true value of *λ* is near zero. The non-zero counts force *γ* ≈ 0 and the BZ model reduces to the base model. This explains why the BZ model performs identically to the base model.

This simulation may be unrealistic, as it is unclear what experiment would cause a random but complete inability to measure a transcript within only select samples in a batch (sample-specific). So we simulated a second dataset of batch-specific complete technical zeros. In simulation 4, a single transcript is measured in 15 replicate samples: 5 replicates in each of 3 batches. However, due to the use of a different reagent or a missed experimental step, within batch 2 there is a complete lack of the transcript. We assume that no other bias is present in batches 1 or 3, which are represented as random draws from a Poisson distribution with rate parameter 1. The results appear similar to those of simulation 3. The difference is that the RI model performs better than the base or BZ models but still underestimates the true value of *λ*. The ZIP model slightly overestimates *λ*. These results of the RI and ZIP models stem from each model’s inability to distinguish between which zeros are due to a sampling process and which are due to a technical process. The ZIP model performs well only in a subset of complete technical processes, *e.g.*, simulation 3, but may still cause over-inflation of parameter estimates in other complete technical processes (*e.g.*, simulation 4).

#### 1.4 Simulation 5: Highlighting Biological Zeros

The fifth simulation consists of 15 samples from three individuals with Poisson rate parameters 1.4, 0, and 3.2. This simulates a situation where the abundance of a single transcript is measured in three individuals: two possess that transcript and one does not. As in the previous simulations, the small rate parameters ensure that this simulation contains sampling zeros as well as biological zeros. To simulate a situation in which the ZIP model is used in a condition-specific way, we modify the ZIP model by replacing 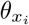 with 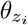. This changes modeling zero-inflation by batch to modeling zero-inflation by individual. We summarize and provide an intuitive explanation of the results (shown in Figure 2E and S7):

##### PC model

The PC model performs poorly, providing biased estimates in all three people^2^.

##### Base model

The base model performs well in this simulation. With no non-zero counts in person 2, the base model places posterior estimates of *λ*_2_ on low values that would be expected to produce large numbers of sampling zeros.

##### ZIP model

The ZIP model massively overestimates value of *λ*_2_ which was so high that the posterior credible intervals were cropped in 2E to aid visualization of the other results. This behavior of the ZIP model comes from the same mechanism that inflated parameter estimates in simulations 1, 2 and 4. Namely, the ZIP model has difficulty distinguishing between high abundance and high zero inflation (high *λ*_2_ and high *θ*_2_) and low abundance and low zero inflation (low *θ*_2_ and low *λ*_2_). The difficulty is far more severe, as all replicates from person 2 are zero and thus the ZIP model has no information to identify this model. This conclusion is supported by Figure S8 which demonstrates how the regions of highest posterior probability span both very high and very low values of *θ*_2_ as the values of *λ*_2_ vary over nearly 10 orders of magnitude.

##### BZ model

The BZ model performs well in this simulation and estimates *λ* well in all 3 people. To see the differences between the base and BZ model results, the estimates for *λ*_2_ are shown on a log scale in Figure S7. The complication of biological zeros is emphasized as on a log scale, the true value of *λ*_2_ is negative infinity. Neither model can estimate this true value due to numerical precision limitations of computers and our use of HMCMC, which cannot handle a latent Dirac distribution and requires an approximating truncated normal distribution (*Methods*). But the zero inflation in the BZ model estimates values of *λ*_2_ approximately two orders of magnitude smaller than the base model. The BZ model places significant posterior probability on large values of *γ*_2_ which also gives this posterior estimate a distinctive bimodal shape. If we had inferred the BZ model with an algorithm that included a latent Dirac distribution, such as a Metropolis-within-Gibbs sampling scheme, the BZ model might place non-negligible probability mass exactly on *λ*_2_ = 0.

## 2 Non-Condition-Specific Zero-Inflation

To investigate whether our results using the ZINB model were unique to condition-specific zero-inflation models, we repeated our analysis with a ZINB model that was fixed to only infer non-condition-specific zero-inflation (see Section 6 for more details). While we find less discrepancy between the NB and non-condition-specific ZINB model (ncsZINB), the observed patterns are similar with an average discrepancy of 32.7% (range: 3.0%-48.0%) among the top-50 most differentially expressed sequences. Similarly, for the top-5 most differentially expressed sequences the average disagreement averaged 23.0% and reached 60.0% for one dataset (Figures S3-S5). In parallel to the condition-specific case, we again observe a strong correlation between the difference in inferred differential expression between the ncsZINB and NB models and the inferred zero-inflation (absolute value of Spearman rho > 0.08 and p-value ≈ 0 for all 6 datasets; Figure S3). That is, even in the absence of condition-specific zero-inflation, the ncsZINB model interpreted that zeros in presence-absence-like cases were evidence of high levels of zero inflation rather than evidence of differential expression.

**Figure S1:**
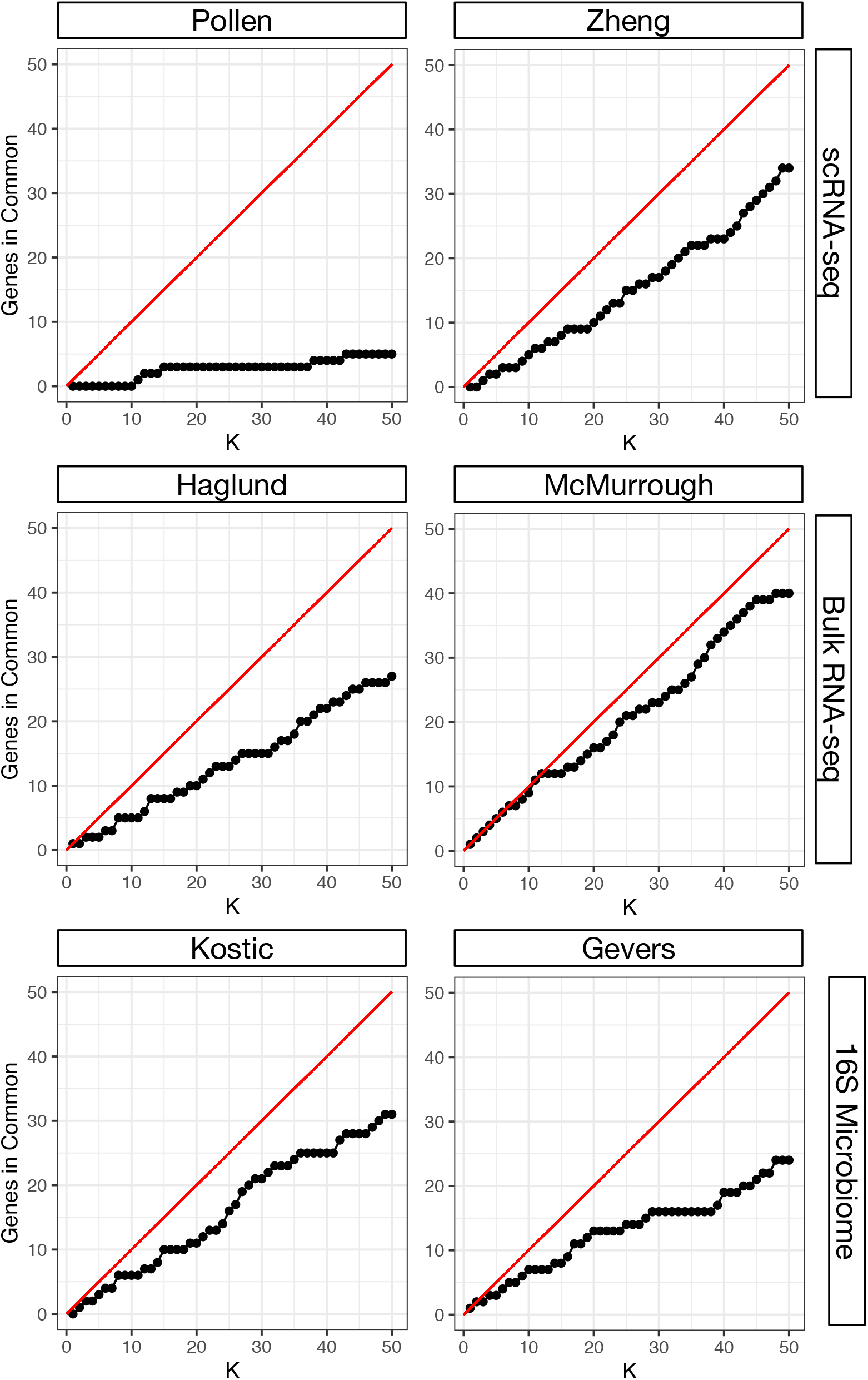
The ZINB and NB models often disagree regarding which sequences are the most differential expressed. For each dataset the intersection between the top-K most deferentially expressed sequences according to the NB and ZINB models is shown as a function of K.

**Figure S2:**
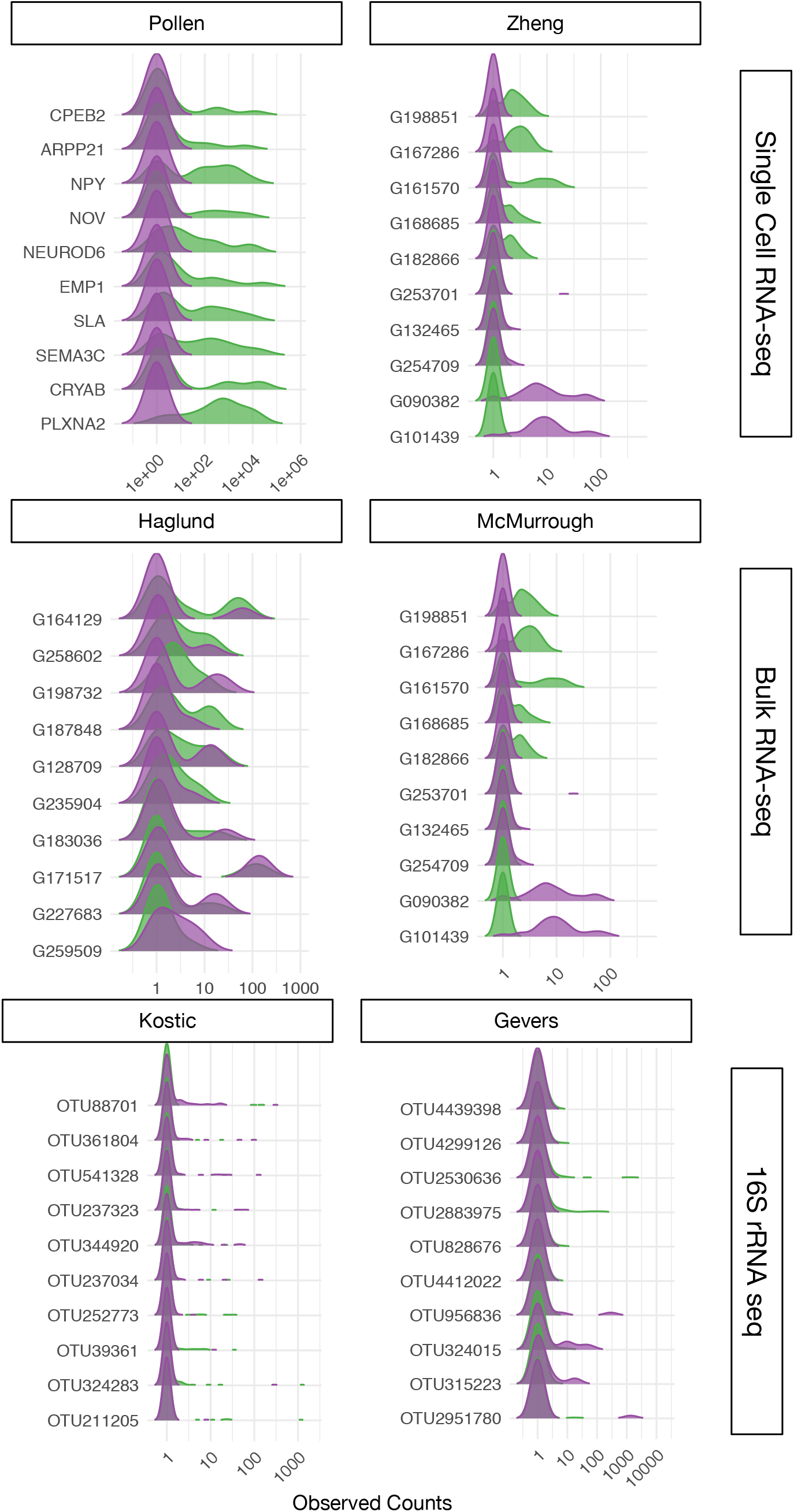
Kernel Density Estimates of observed count values in each condition for the 10 sequences from Figure 1 that had the largest discrepancy in their estimated differential expression between the NB and ZINB models.

**Figure S3:**
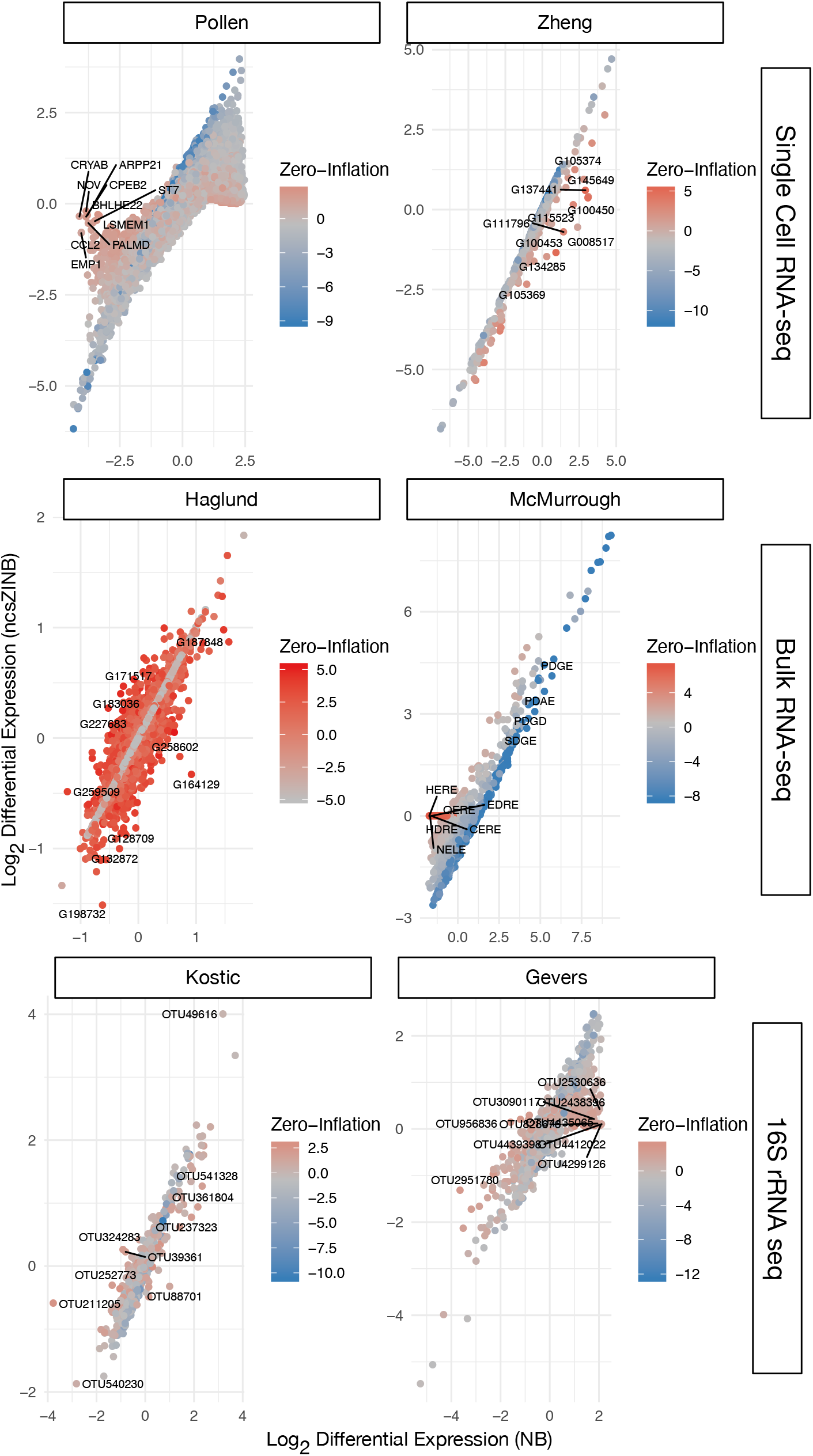
Differential expression (DE) estimates from a negative binomial (NB) and non-condition-specific zero-inflated negative binomial (ncsZINB) model can differ substantially. Log base 2 differential expression for the ncsZINB and NB models are shown after each was applied to single cell RNA-seq, bulk RNA-seq, and 16S rRNA microbiota data. Dots represent different sequences, and each is colored according to the degree of zero-inflation as estimated by the ncsZINB model. For each dataset, the 10 sequences that have the largest discrepancy between inferred DE are labeled and their distribution in each condition is given in Figure S5.

**Figure S4:**
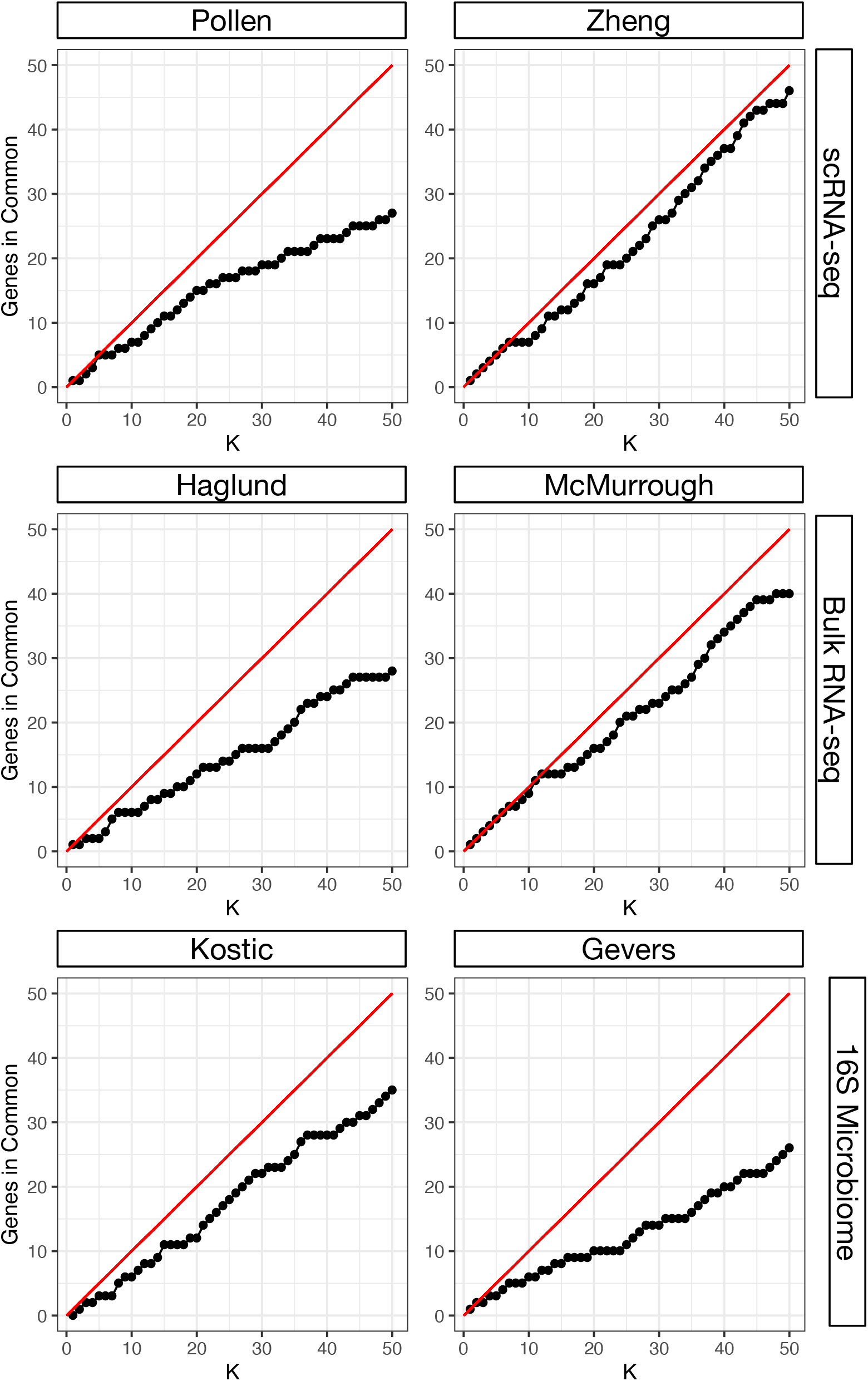
The ncsZINB and NB models often disagree regarding which sequences are the most differential expressed. For each dataset the intersection between the top-K most deferentially expressed sequences according to the NB and ncsZINB models is shown as a function of K.

**Figure S5:**
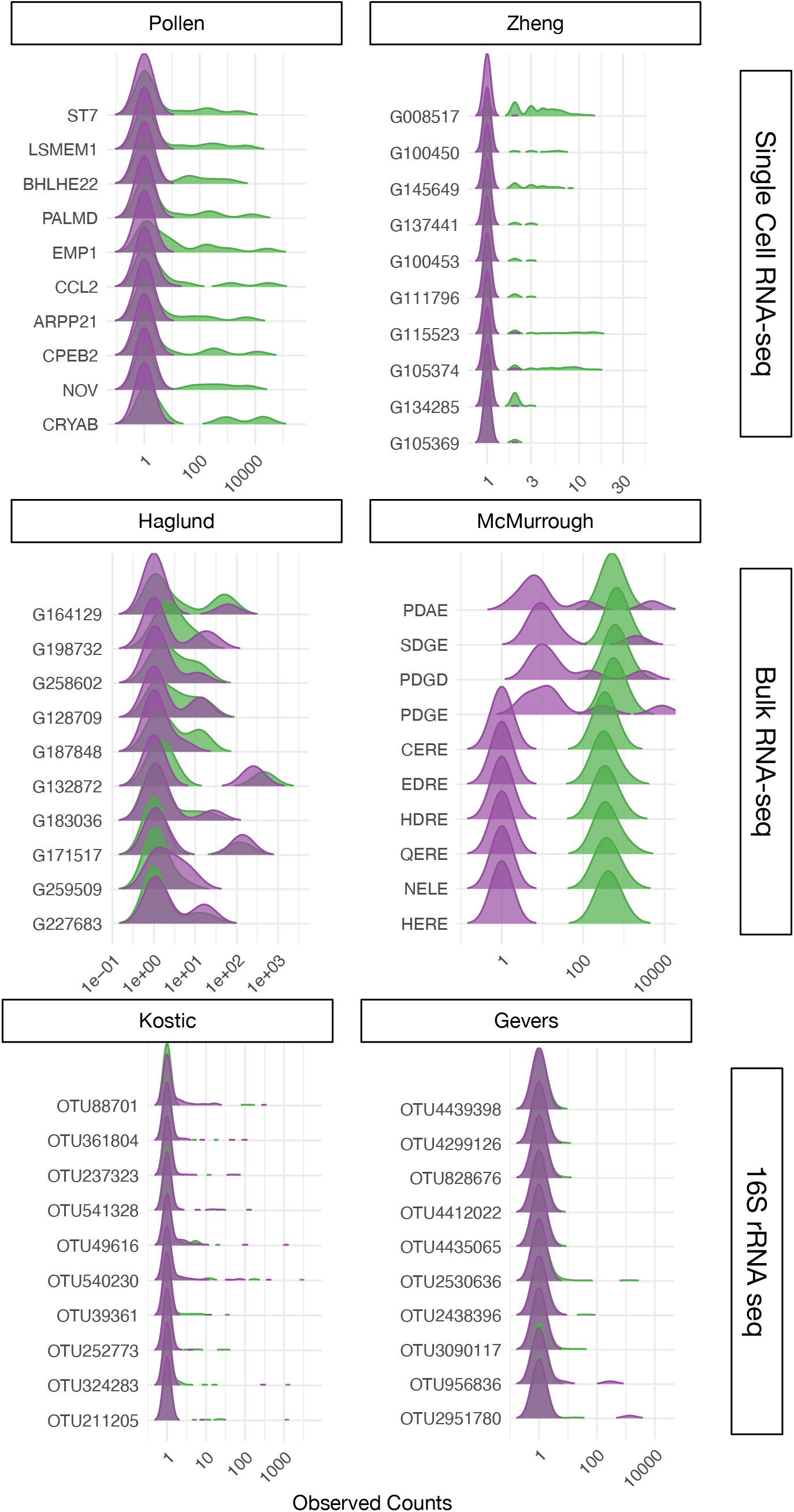
Kernel Density Estimates of observed count values in each condition for the 10 sequences in Figure S3 that had the largest discrepancy in their estimated differential expression between the NB and ncsZINB models.

**Figure S6:**
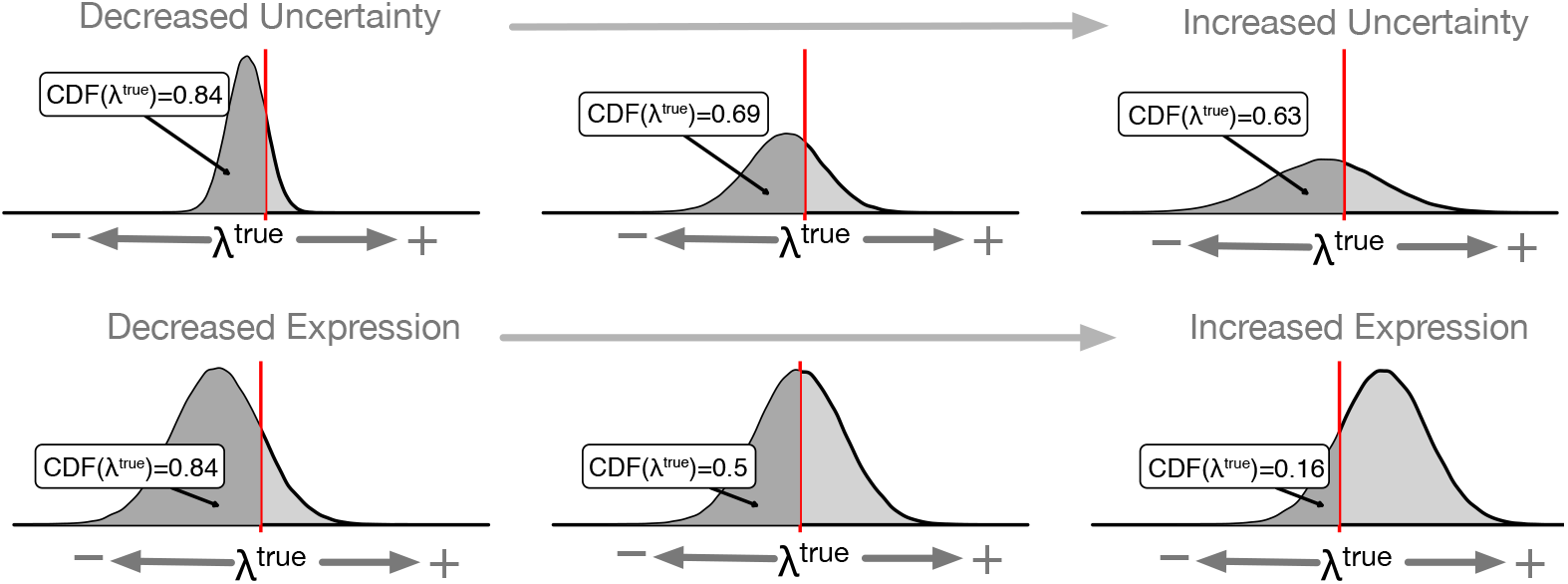
The cumulative distribution function (CDF) of the posterior distribution is a function that for any value of *λ* calculates the integral of the density from negative infinity to the specified value of *λ*. If we set that value of *λ* to be equal to the true value of *λ* in the simulation, then *CDF* (*λ^true^*) is a measure of how close the mean of the density is to the true value as well as accounting for how diffuse the density is about the true value. When each density represents the posterior distribution of a model then this statistic makes a suitable performance measure for how accurately and how precisely a model inferred the true value of *λ*. An optimal model will produce a posterior distribution where *CDF* (*λ^true^*) = 0.5.

**Figure S7:**
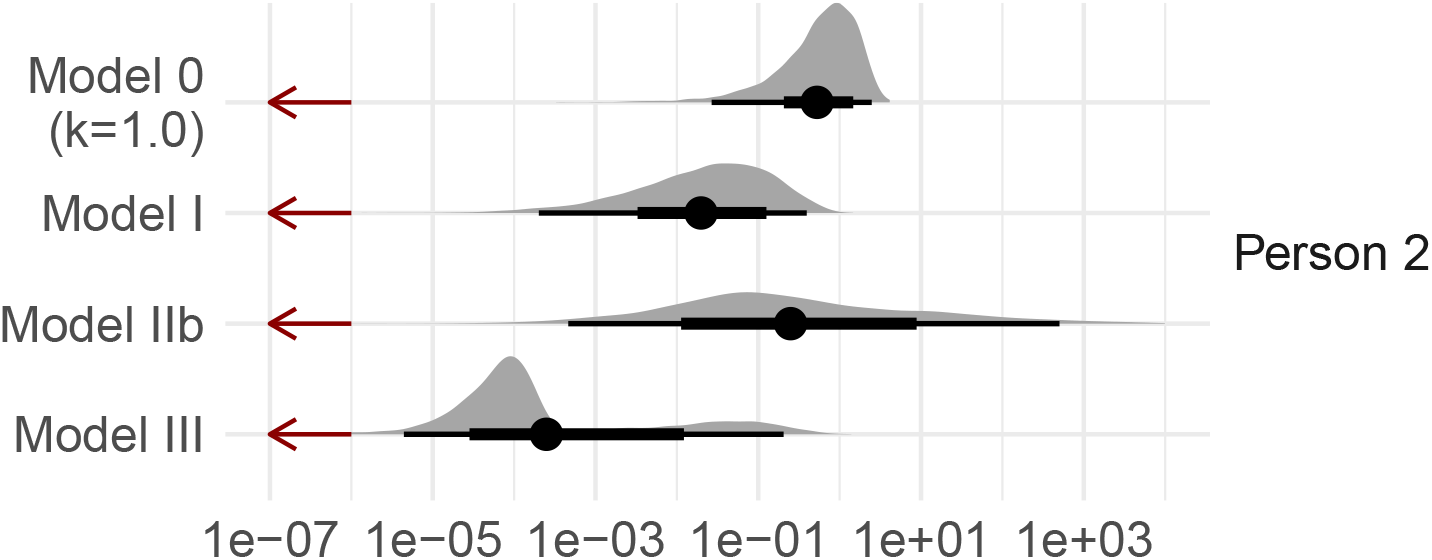
Posterior distribution of *λ* from each model applied to Simulation 5 shown on a log-scale for Person 2–an example of biological zeros and sampling zeros. Dark red vertical bar represents the true value of *λ*. Posterior mean as well as the 66% and 95% credible intervals are shown in black.

**Figure S8:**
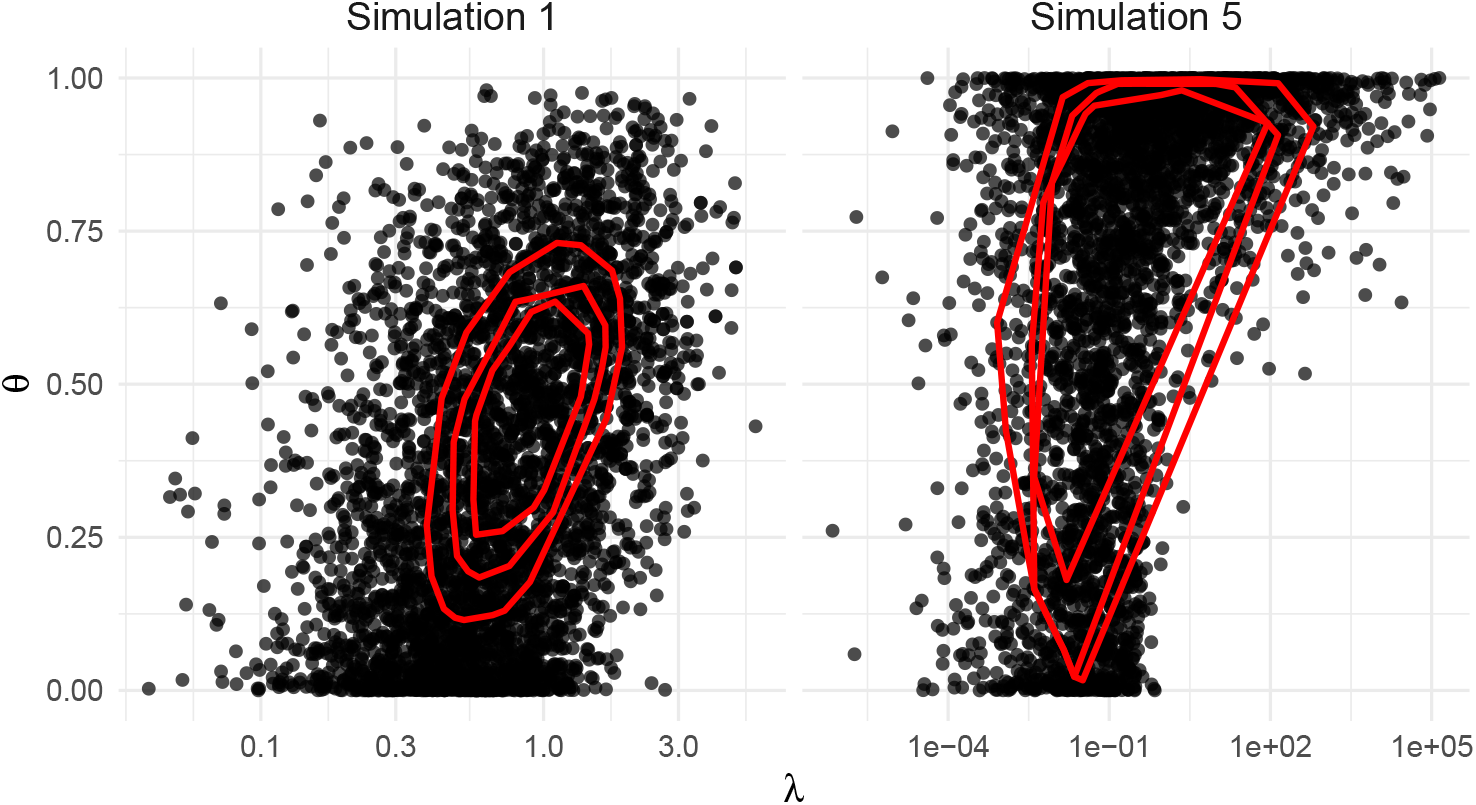
Large uncertainty explains the parameter inflation observed with the ZIP model. Posterior samples of *λ* (transcript abundance) and *θ* (probability of zero inflation) for the ZIP model applied to simulation 1 (sampling zeros) and simulation 5 (biological zeros). For simulation 5, the posterior distribution is of *λ*_2_ and *θ*_2_. The ZIP model is unable to distinguish between zeros due to sampling (i.e., low *λ* and low *θ*) versus zeros due to zero inflation (i.e., high *θ* and either low or high *λ*). Note that for Simulation 5, this uncertainty over *λ*_2_ spans nearly 10 orders of magnitude. The 80%, 90%, and 95% highest posterior density regions for the log posterior probability are shown in red.

**Figure S9:**
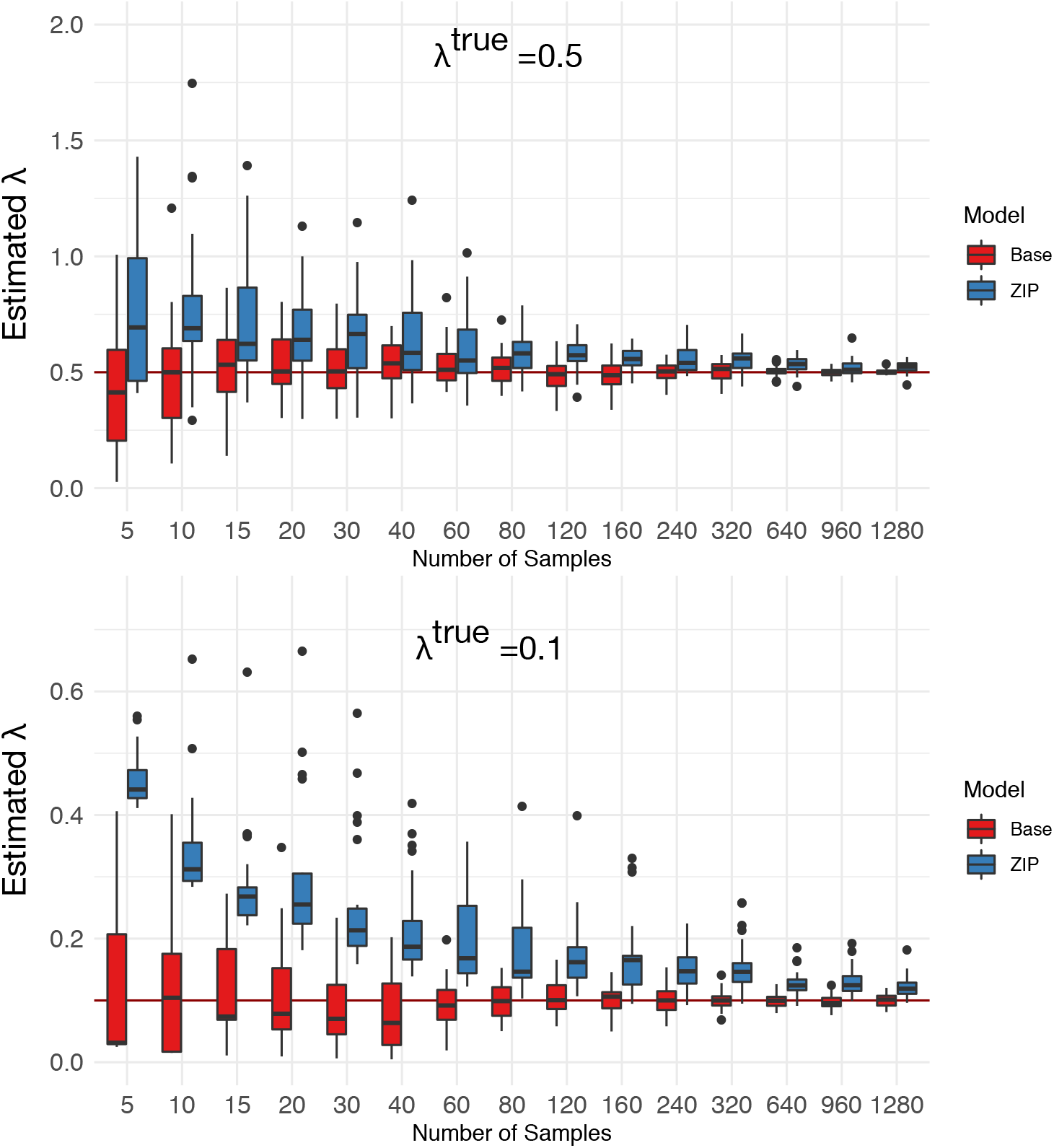
With sample sizes between 5 and 1280, 30 datasets were simulated and analyzed with the Base and ZIP models. Each simulated dataset contained only sampling zeros (simulated with Poisson rate (*λ*) parameter of 0.5 - top panel and 0.1 - bottom panel). Estimates from the ZIP model show substantial bias even for sample sizes larger than 1000. This bias increases, and takes more samples to mitigate as *λ^true^* decreases.

**Figure S10:**
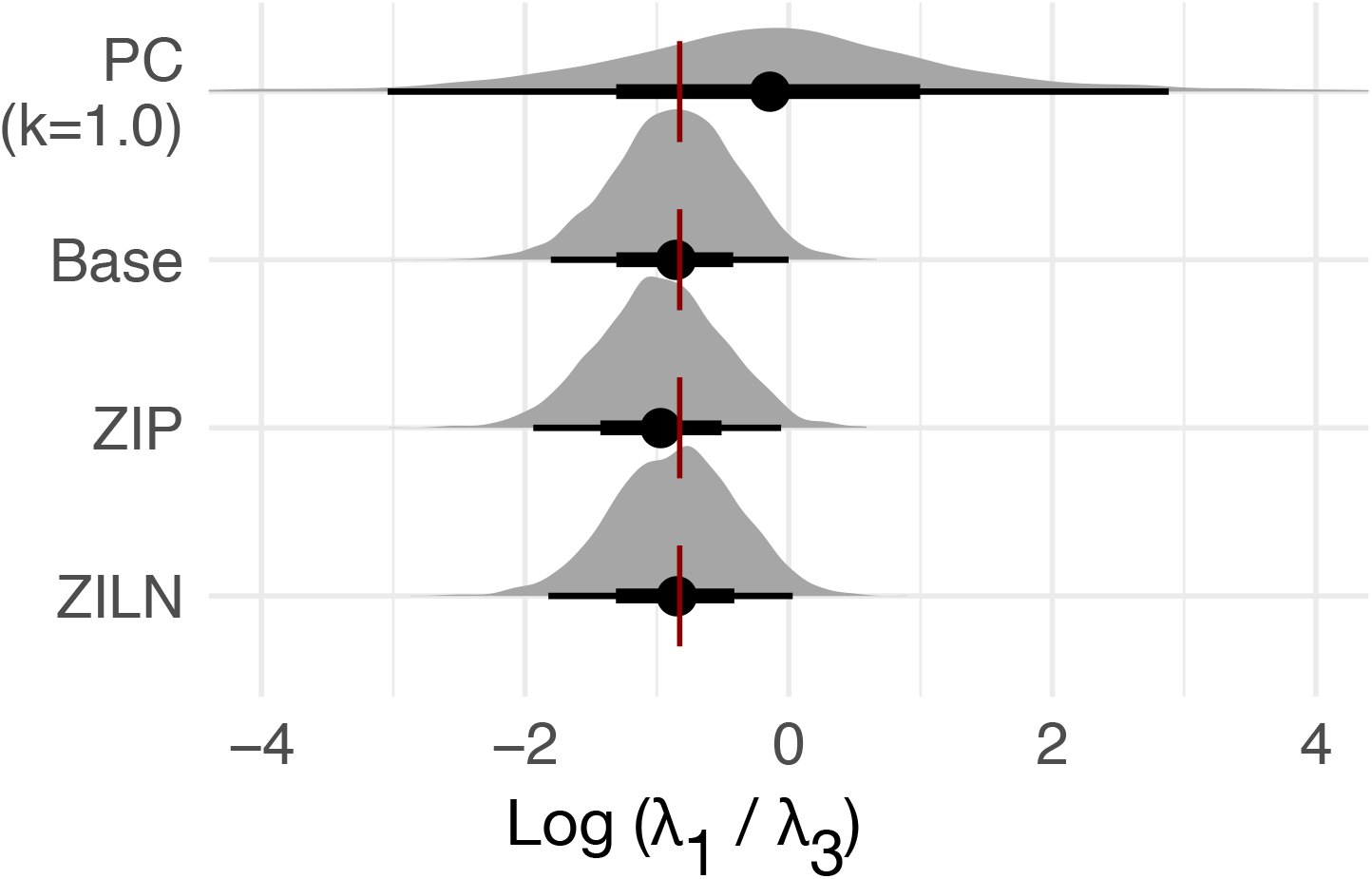
Posterior distribution of the differential expression 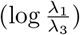 of a simulated transcript between persons 1 and 3 in Simulation 5. Dark red vertical bar represents true value of *λ*. Posterior mean as well as the 66% and 95% credible intervals are shown in black.

---

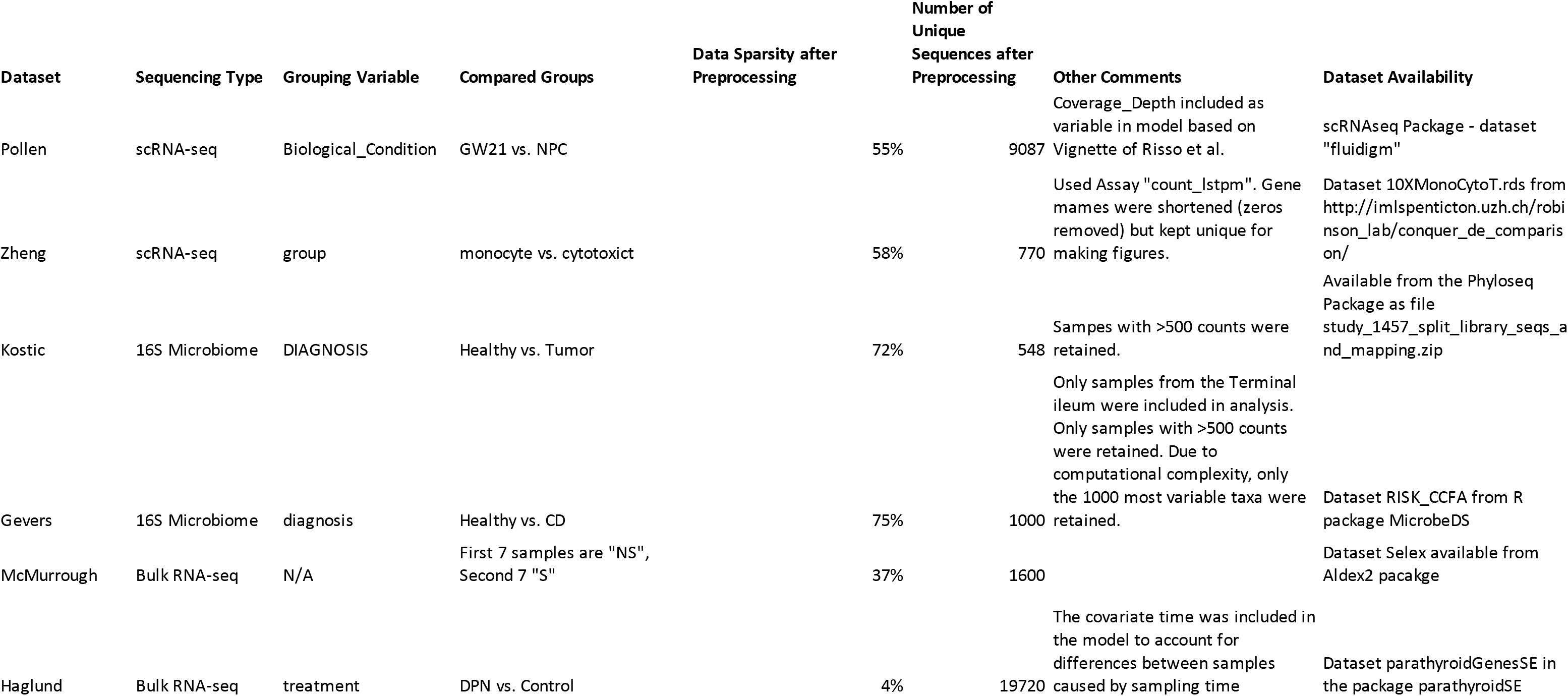

## Footnotes

1 That the ZIP model’s biased estimates improve with increasing sample size at all is because the model uses the variation of the non-zero counts to eventually approach the correct answer.

2 We included the PC model to show how including a fixed pseudo-count forces the posterior estimates for *λ*_2_ to remain near the pseudo-count value, without allowing the model to approach the true value of *λ*_2_ = 0.

## Notes

#### Summary of Updates

Our real data analysis now features an analysis of 5 new datasets, including data from 16S rRNA microbiome studies and bulk RNA-seq studies. We have rewritten sections to enhance readability.

